# HiFi-Helper: A reproducible workflow for genome assembly from HiFi reads alone

**DOI:** 10.64898/2026.03.11.710913

**Authors:** Krista Pipho, Greg A. Wray

## Abstract

The PacBio Revio simplifies genome assembly by generating very long reads with very few errors at an affordable price point. Comparative ease of assembly is democratizing access, leading to a larger niche for assembly workflows. HiFi-Helper is a user-friendly snakemake workflow designed to facilitate genome assembly from HiFi data alone. This tool produces a visual summary that provides intuitive feedback on the quality of the resulting assembly and guides assembly parameter optimization. The case studies presented here confirm that HiFi-only assemblies produced by HiFi-helper can meet or exceed the quality of genomes assembled in prior decades with a much lower investment of time and resources.

## 1 Introduction

The PacBio Revio changes the genome assembly game by generating very long reads with very few errors at an affordable price point (Hon et al., 2020). This innovation marks the dawn of ‘easy’ non-model organism reference genome assembly. Comparative ease of assembly is democratizing access, meaning that labs without dedicated bioinformaticians and massive sequencing budgets are undertaking assembly projects in higher numbers. These projects will need assembly code designed for PacBio HiFi reads that is user-friendly, has the option to accommodate minimal computing resources, and provides intuitive feedback on the quality of the resulting assembly.

Colora is one published option that fulfills many of these requirements, and can create phased chromosome level assemblies from high quality datasets (Obinu et al., 2025). One drawback is that the workflow requires Hi-C reads to use in assembly. For many small diploid genomes, HiFi data alone can create a respectable first assembly or an improvement over existing assemblies made in preceding decades. Thus there is a gap to be filled in terms of feature-rich assembly workflows that support HiFi-only assembly. Given this, we offer HiFi-Helper as a simple, user-friendly tool for assembling HiFi reads.

HiFi-Helper is built in Snakemake, a python based bioinformatics workflow language offering modular analysis optimized for both personal and high performance computing settings (Koster and Rahmann, 2012). To manage the software packages and tools needed for execution of the workflow we utilize pixi. Pixi is built around the conda ecosystem, but it runs faster, provides a lock file for improved reproducibility, and stores environments in the relevant local directory rather than in a distant part of the file system (prefix-dev, 2026). Together, Snakemake and Pixi make HiFi-Helper easy to install and run.

The workflow supports two main types of usage. 1) Processing PacBio HiFi reads into a de novo assembly and providing quality metrics compared to a provided reference genome or 2) Taking two assembled genomes and providing quality metrics comparing them. In both cases R is used to synthesize multiple core output files into an intuitive visual summary that gives an easy overview of assembly quality. This means that a ‘reference’ genome is always required as input despite this reference not being used during assembly. In practice, these two modes can be used iteratively, using multiple rounds of assembly parameter optimization and output comparison to arrive at the best genome possible from limited data. This manuscript first offers an overview of tools available in the workflow. Later, under the Results heading, it offers two case studies of how the workflow can be used iteratively to improve final assembly quality.

## 2 Central Tools

PacBio’s own HiFiasm software is at the heart of the workflow. HiFiasm comes with the expected perks of an enterprise-level development team; it is very efficient and incredibly well documented. In HiFi-Helper the assembly commands are situated in an easy-to-access sub module located at bin/assembly.sh. This means that a user can refer to the hifiasm manual page to customize any part of the assembly command. It is also possible to switch this code out for another assembler entirely if the end user desires.

The only two non-optional analyses are Quast and BUSCO. Quast is a lightweight software originally designed for benchmarking the output of different assembly programs against each other (Gurevich et al., 2013). It quantifies the final number of un-connected sequences (or “contigs”) produced by an assembler, along with the range of contig sizes and how many of the largest pieces it takes to make up a given % of the total sequence length (LC50 for 50%, LC75 for 75%). BUSCO, an abbreviation for Benchmarking Universal Single-Copy Orthologs, is a program which assesses the completeness of genome assemblies by comparison to a theoretical biological expectation of their content (Simao et al., 2015). The BUSCO database catalogues known single copy orthologs conserved within different taxa, and the BUSCO software quantifies the number of these biologically expected sequences that are captured in an assembly.

In HiFi-Helper, the use of BUSCO is two-fold. First and foremost the program is used for its intended purpose of checking assembly quality by confirming the presence of conserved single-copy orthologs. Secondarily, the BUSCO outputs already generated for checking completeness are re-purposed as a quick way to identify homologous chromosomes between the genome of interest and a comparison assembly provided for analysis.

During development of this workflow test data from yeast was used as a quick way to run and troubleshoot the analysis. The sample provided for analysis is SRR13577847, a small and easily downloadable PacBio HiFi sequencing dataset from an S288c-like yeast strain (SRA, 2021). The comparison assembly used alongside this sample is the S288c reference genome from the NCBI genome database (NCBI, 2014). These two assets were included as the default settings on download, so that users can quickly confirm that the workflow executes without errors in their computing environment.

Finding and interpreting the outputs of large workflows can sometimes be a substantial task. HiFi-Helper uses two main strategies to make outputs more accessible. First, while a large number of important final outputs are automatically copied into a results folder, the two most important outputs, genome FASTAs and visual summaries, are also placed in the outermost folder that the user interacts with directly. Second, this visual summary produced using R-Markdown provides an HTML page displaying critical information from the core workflow analyses in a centralized and easy to interpret format (“R-markdown”, 2026).

## 3 Optional Capabilities

In addition to the core genome assembly and comparison functionality of HiFi-helper, there are several optional utility features. Pre-assembly utilities include HiFi read contaminant filtering, kmer based HiFi read quality modeling, and targeted mitochondrial genome assembly from HiFi reads. Post assembly utilities feature kmer based comparison of two genome FASTAs, telomere finding from a genome FASTA, and repeat masking of a genome FASTA. Each optional feature is described in more detail below.

Pre-Assembly:

The HiFi read contaminant filtering is performed with Kraken 2 (Wood et al., 2019). The creators of Kraken2 provide a selection of databases consisting of known sequences, including bacteria, fungi, and common research species. A kmer based alignment approach is then used to search input reads for matches to known sequences. These matches are later used to categorize HiFi reads by species of origin. This is a very useful way to remove contaminating reads from the sample’s microbiome and infections, or from laboratory contact with bacteria, fungi, other research samples, or the human experimenter’s own tissue. This workflow assumes that the sample species in question is not present in the Kraken2 database, and thus removes all reads assigned a species category from the input. A note is made in the README file to modify this behavior if the user is working with a common research species or a bacterial or fungal sample.

HiFi read quality modeling is done using Genomescope2 (Ranallo-Benavidez et al., 2020). Genomescope2 is a fast and simple but statistically sophisticated approach to examining reads without alignment to a reference genome. The program counts the instances of all possible short sequences, or kmers, within the input reads to create a so-called kmer spectrum. The spectrum is then used to construct mathematical models of the genome giving rise to the input reads. This model estimates parameters such as genome size, ploidy and heterozygosity. In addition it provides an estimate of sequencing error rates, sequencing coverage, and (based on self report of model confidence) a quick and general sanity check of whether the sequencing data is sufficient for the desired downstream research task.

Targeted mitochondrial genome assembly is undertaken using the Oatk package and database (Zhou et al., 2025). Although originally developed for the complex assembly task of sorting out the genomes of plant mitochondria and chloroplasts, the database contains organelle genome sequences from a wide range of taxa including many animals. In addition to the edge conferred by taxonomically relevant reference sequence databases, the Oatk assembler is optimized for detection of coverage imbalances between organelle genomes and the nuclear genome. For our purposes, mitochondrial reads are often present at a substantially higher coverage than the nuclear genome, which helps to disentangle lateral sequence transfer events to the nuclear genome from true mitochondrial sequence.

Post-Assembly:

In terms of post-assembly features, one of the most broadly useful optional tools of HiFi-Helper is the sample genome x comparison genome dotplot. Appropriate outputs are generated for use with Maria Nattestad’s online interactive dotplot viewer (Nattestad, 2017), which allows exploration of a detailed alignment between the two genome FASTAs. Features include minimal initial data loading for fast interactivity, plot color selection, and interactive zoom. The tool facilitates intuitive qualitative assessment of synteny, chromosome structure, repeat content, and sequence divergence. The underlying data is generated using nucmer, a central offering of the Mummer4 analysis package (Marcais et al., 2018). Nucmer uses a kmer based technique to align large and potentially highly diverged sequences, and provides ideal output data for visualizing repeated or inverted regions.

A simple strategy often employed as a first check of chromosome completeness in an assembly is the presence of telomeric sequences on both ends. HiFi-helper offers an optional telomere finding step using the Telomere Identification Toolkit, or TIDK (Brown et al., 2025). This tool creates a simple visual output that shows the location and density of telomeric repeats, and can easily be used to tell by eye if both ends of a scaffold have substantial telomeric content. TIDK also offers a telomeric sequence database to easily look up which sequence should be input for your organism of interest’s telomeric repeat.

Repeat masking of a reference genome is an important step to take prior to many types of downstream analysis. For example, masking repetitive regions prior to aligning reads can lead to faster and less noisy analysis. Masking with a tool like RepeatMasker is a fast and simple process if the sequences to mask are already identified (Tarailo-Graovac et al., 2009). For a newly assembled genome though, especially one that is evolutionarily distant from well-studied taxa, existing repeat databases may not contain many of the highly repetitive sequences truly present in the new genome. In this case a computationally expensive tool like RepeatModeler is beneficial to use first (Flynn et al., 2020). This software builds a new database of repetitive sequences tailored specifically to the genome in question. The output of this tool can then be used to run RepeatMasker. HiFi-helper supports repeat modeling for the sample genome, or providing an existing repeat database for substantial compute-time savings.

## 4 Case Study

### 4.1 Heliconius melpomene rosina

The Heliconius melpomene rosina (HMR) genome is relatively small and repeat poor compared to other members of the genus, which made it an organism of choice for early assembly projects. Several rounds of effort across assembly versions Hmel1, Hmel2 and Hmel2.5 have yielded a relatively high quality assembly informed by long read sequencing and linkage data. New PacBio HiFi data was collected from a female individual in an attempt to capture, for the first time in a Heliconius reference genome, the W chromosome. The assembly process for this project is used here as an example of the ideal HiFi-only assembly situation with sufficient HiFi read coverage and a highly useful existing reference genome.

A HiFi-Helper run with default settings was run on HMR x Hmel2.5. Plots from the visual summary are shown and interpretation is discussed. From the content of Figure 1 we learn that generating an HMR assembly using the HiFiasm default parameters attains a surprisingly comparable assembly quality to the existing best in genus reference. Figure 2 shows the shared BUSCO heatmap, which is a tool for visualizing shared sequence between contigs from two different assemblies. The shared BUSCO counts are log2 transformed to facilitate visualization of small BUSCO overlap values. This simple visual highlights the shortcomings of HiFi-only assembly with identification of a large structural error in HMR relative to the Hmel2.5 assembly constructed using linkage group information.

**Figure 1:**
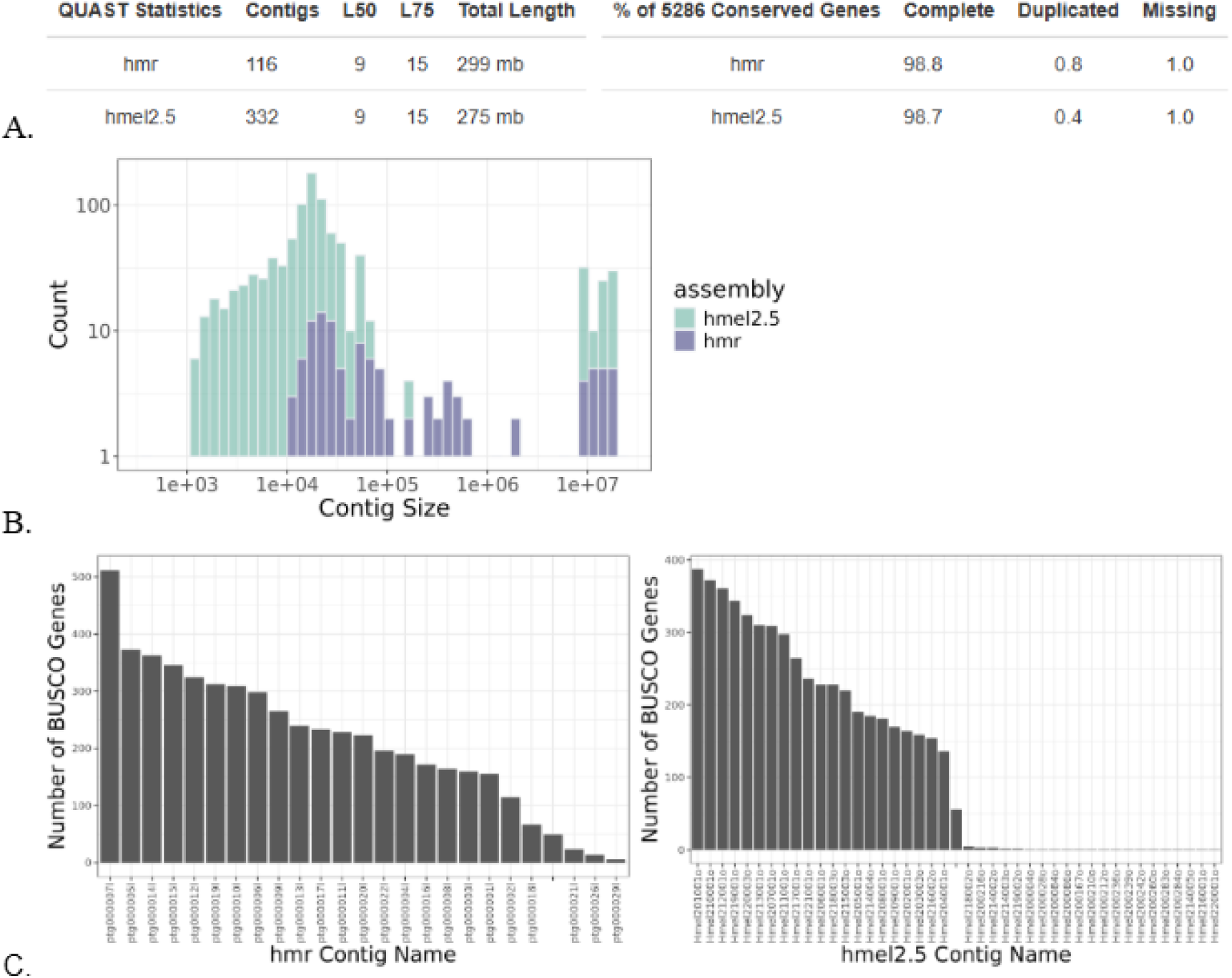
Assessment of HMR Assembly Quality Using the Visual Summary. a) The overview table shows that the default-settings HMR assembly has fewer contigs, slightly larger overall length, and a modest increase in BUSCO gene model duplication rate, but is otherwise highly comparable to Hmel2.5. b) The first graph in the visual summary shows the distribution of contig sizes pulled from the assembly fasta.fai index files. In this case it shows similar contig size distributions for the two assemblies, with HMR having a very similar set of chromosome sized contigs but fewer and longer minor contigs. c) The second and third graphs that appear in the visual summary show the number of BUSCO genes found on each contig. In the HMR assembly 21 major contigs with a large number of BUSCOS corresponds to the known 21 chromosomes of this species. The most salient difference from Hmel2.5 is that the default-setting HMR assembly has fewer BUSCO gene models stranded on minor contigs.

**Figure 2:**
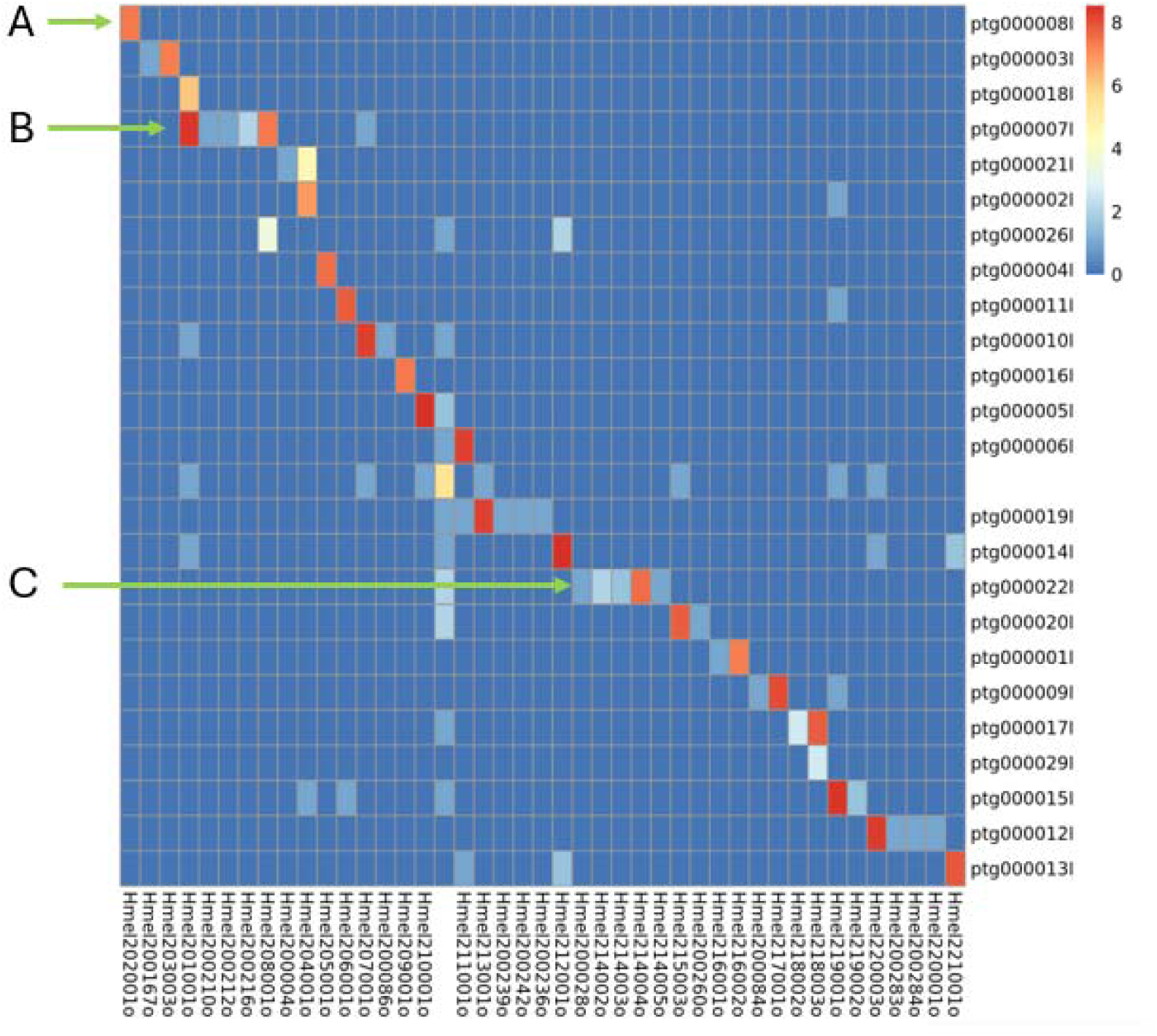
Assessment of Chromosome-Level Homology Using the Visual Summary. a) This orange square shows high overlap in found BUSCO genes between Hmel202001o (Chromosome 2) and HMR contig ptg000008l. The dark blue on this vertical shows that Hmel202001o does not share identified BUSCO genes with any other HMR contigs. The dark blue on this horizontal shows that ptg000008l does not share identified BUSCO genes with any other Hmel2.5 contig.Thus the assignment of homology between these two contigs is unambiguous. b) The cluster of this red square, the light orange square above it, and the orange square four spaces to the right signals a potential problem with the default HMR assembly. HMR contig ptg000007l shares substantial BUSCO gene overlap with two Hmel2.5 contigs (Hmel201001o and Hmel208001o), while Hmel2.5 contig Hmel201001o shares substantial BUSCOS with two HMR contigs (ptg000007l and ptg000018l). This suggests a major and complex structural difference between HMR and Hmel2.5. Because Hmel2.5 incorporates linkage data and HMR does not, this is presumed to be a miss-assembly in HMR. c) On the other hand, HMR shows several cases of successfully joining sequences known by linkage to be on the same chromosome. Three of the four pale squares in this area represent the small number of BUSCOS on minor Hmel2.5 contigs Hmel214002o, Hmel214003o and Hmel214005o being joined with the major contig Hmel214004o into a more complete sequence of Chromosome 14 in HMR ptg000022l.

**Figure 3:**
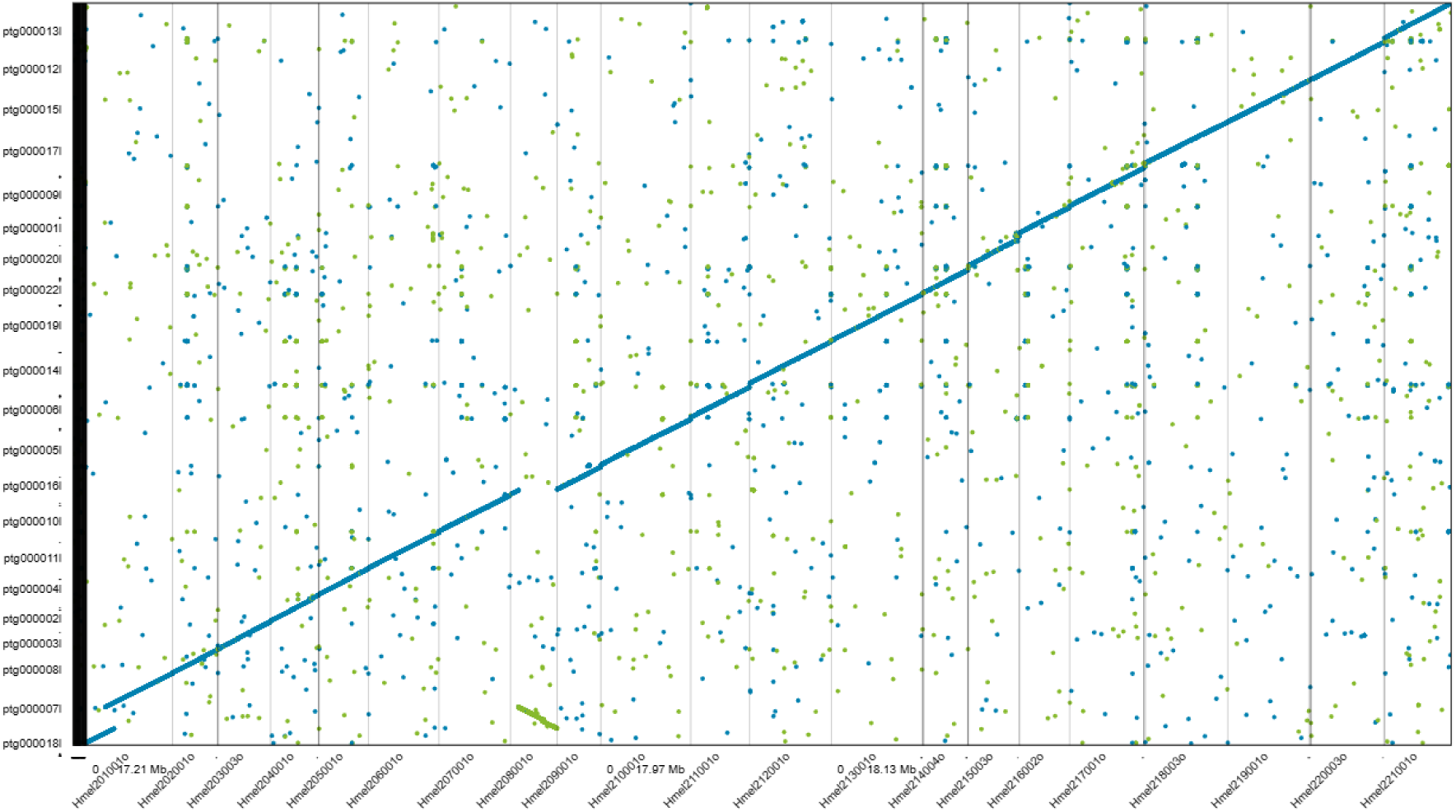
Major Structural Error Visible on Interactive Dotplot. Mummer4-nucmer was used to align the Hmel2.5 genome (x axis) to the HMR assembly (y axis). A kmer size of 20 was used for this analysis. Unique forward kmer overlaps are shown in blue and unique reverse kmer overlaps are shown in green. This visualization method confirms the structural issue with assembly of chromosome one and makes apparent that Chromosome 8 has been erroneously assembled into the middle of chromosome 1 in the reverse orientation.

If we are willing to put more computational resources into seeing this more clearly we can turn on the optional generate_data_for_dotplot feature and re-run the workflow. All other expected outputs will already be present, so the Snakemake rules corresponding to them will not re-run. The only rule that should run this time is “dotplot”, which will generate a .coords and .coords.idx file in the results directory. These output files can be visualized using the Dot web tool linked in the config.yaml. Using the web viewer we see that while there is beautiful synteny between hmel2.5 and HMR throughout the rest of the genome, the default HMR assembly has placed almost all of Chromosome 8 in the middle of Chromosome 1.

With this knowledge in hand we can try to improve upon our default assembly of HMR by tailoring HiFiasm to what we know about our data. The genomescope2 kmer spectrum and model from our first run (Figure 4) can tell us several useful things. First, the model estimates the total length of the genome that generated this kmer spectrum, and this can be entered into HiFiasm under the –hg-size parameter. Second, it tells us that the baseline coverage per kmer is 15.3. Looking at the kmer spectrum we can indeed see that there is a leftmost peak at the x axis coverage value of 15, and we know this corresponds to sequences that appear only once in our diploid genome. Further to the right there is a peak at the x axis coverage value of about 30, and we know this corresponds to sequences that appear exactly twice in our diploid genome, or sequences that are identical in both haplotypes. This information can be entered into HiFiasm under the --hom-cov or homozygous coverage parameter. Customizing these two parameters is always an appropriate first optimization step.

**Figure 4:**
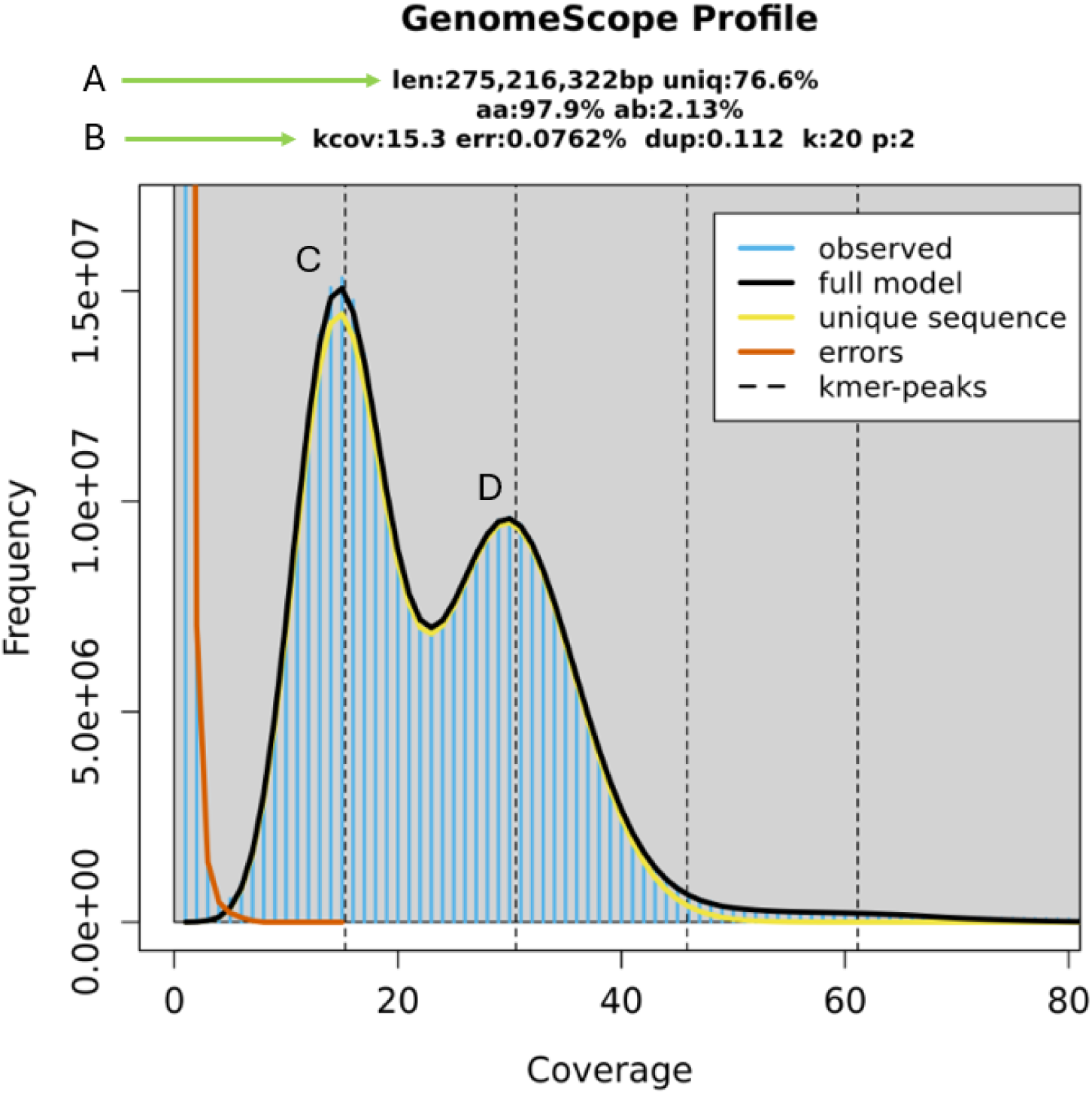
Customizing HiFiasm parameters using Genomescope2. a) The Genomescope2 model calculated total genome length. b) Kmer coverage, or the coverage depth for a single haplotype. c) The peak corresponding to heterozygous sequences, or sequences that only appear in one haplotype. d) The peak corresponding to homozygous sequences, or sequences that appear identically in both haplotypes. Kmers representing higher order repeats are present in the long tail to the right of the graph.

We suspect that the majority of Chromosome 8 was placed incorrectly in HMR due to poor resolution of similar sequences between Hmel201001o and Hmel208001o. To address this the - -max-kocc parameter was increased to 5000 to ensure that even very high incidence kmers are used to resolve ambiguous overlaps. The -N parameter was increased to 300 to ensure that high incidence kmers are included in error correction steps. In addition the TIDK database was used to identify the appropriate telomeric repeat and input it using the –telo-m parameter. In Figure 5 we see that these parameters resolved the misassembly of Chromosome 1 and Chromosome 8, but broke Chromosome 12 into two pieces.

**Figure 5:**
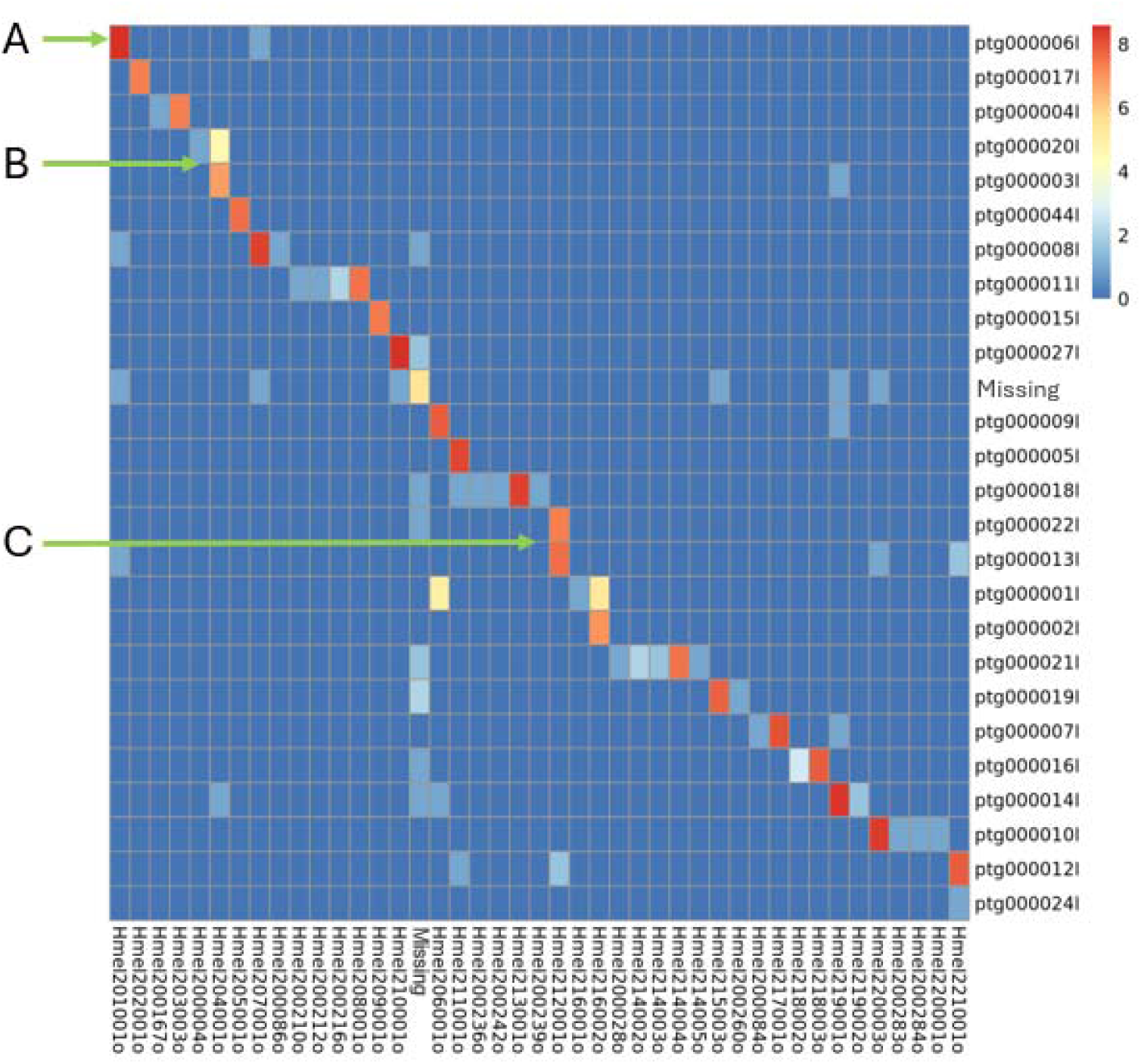
Genome Assembly Parameters Have Trade-offs. a) This bright red square and the dark blue squares below and to the right show that Hmel201001o now has unambiguous homology to ptg000006l. b) The splitting of Hmel204001o BUSCOs between two HMR contigs was present before and has not been resolved based on the new assembly parameters. c) Hmel212001o had an unambiguous homologous contig in the previous assembly, but that scaffold now corresponds to two separate HMR contigs.

These kinds of tradeoffs are common; preventing misassemblies also loses some low confidence connections that were truly present. In this case we can use the linkage information embedded in Hmel2.5 to mitigate this tradeoff. In order to make use of the linkage information in Hmel2.5 we employ the reference guided assembly tool RagTag. Using RagTag’s scaffold feature, our chosen de novo assembled contig set is aligned to Hmel2.5 and contigs aligning to the same reference chromosome are joined in the correct order and orientation with 100 N bases signifying the sequence gaps.

This process yields a new final assembly that makes use of linkage information contained in Hmel2.5. The BUSCO scores are incrementally better, but the major improvement over Hmel2.5 is the placement of all identified BUSCOs into the 21 chromosomal contigs (Figure 6). All major HMR scaffolds correspond unambiguously with the Hmel2.5 linkage groups (Figure 7)(See Appendix B for a shared BUSCO heatmap of .the final assembly), meaning that HMR attains chromosome level scaffolds for all 21 chromosomes.

**Figure 6:**
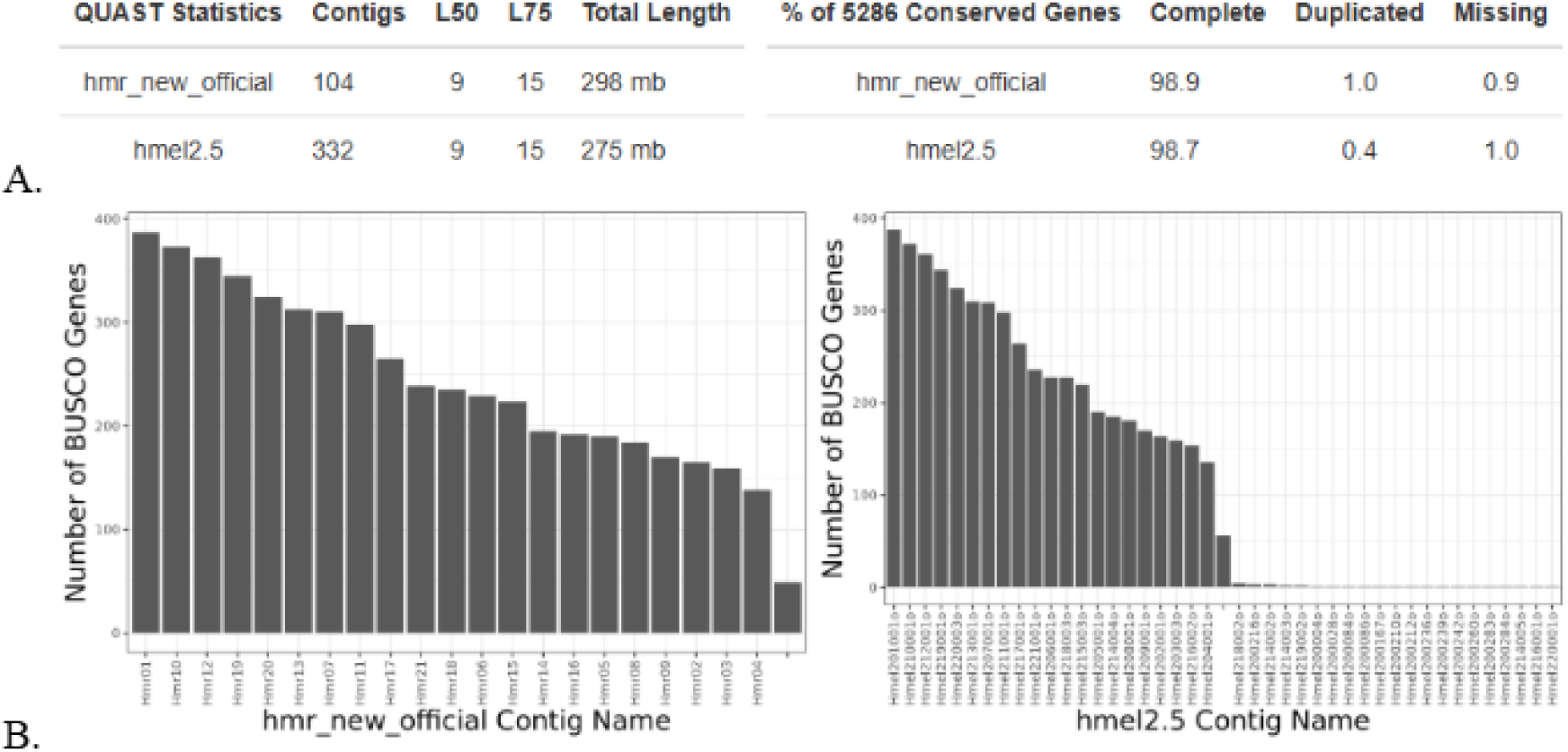
The Final HMR Assembly Improves Placement of Minor Contigs. a) The total genome size of Hmel2.5 is closer to the Genomescope2 genome size estimate, and HMR minorly increases the duplication rate of BUSCO gene models. Both of these factors point to the increased contiguity coming at the cost of introducing a small number of spurious duplications. b) While BUSCO gene models are found across many contigs in Hmel2.5, HMR consolidates all found BUSCOS onto the 21 largest contigs.

**Figure 7:**
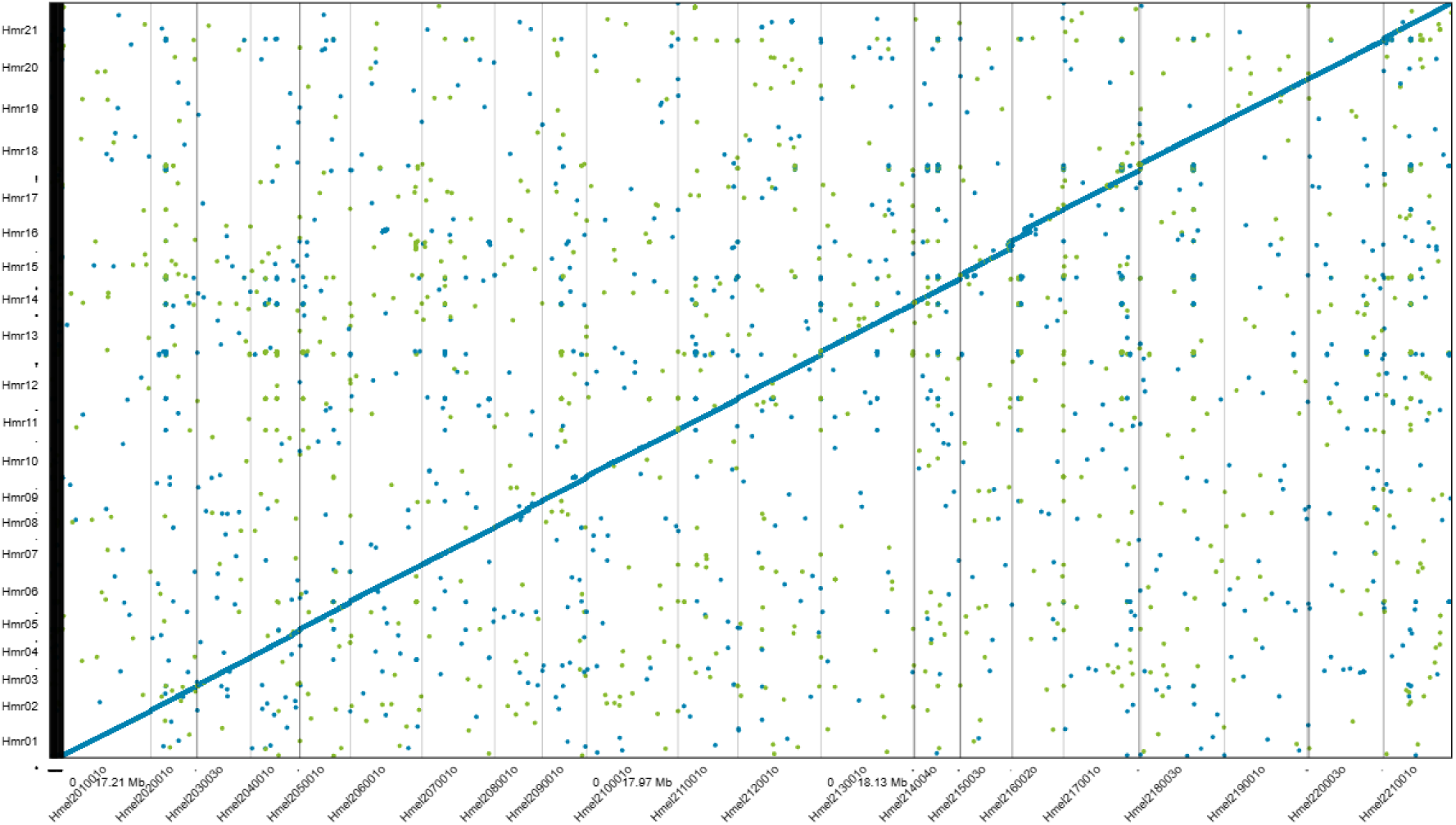
HMR Maintains the Structure of Hmel2.5 While Improving Contiguity.

### 4.2 Dendraster excentricus

At the time of this publication there is one Dendraster excentricus genome present in NCBI, but it contains over 400,000 scaffolds. There is only one other NCBI genome available in the order clypeasteroida, an assembly for the common sand dollar, Echinarachnius parma. Using this as the reference genome and default parameters for HiFiasm we see that the new HiFi reads only assembly is fragmented, but still offers a considerable improvement in genomic information for this part of the tree of life (Figure 8).

**Figure 8:**
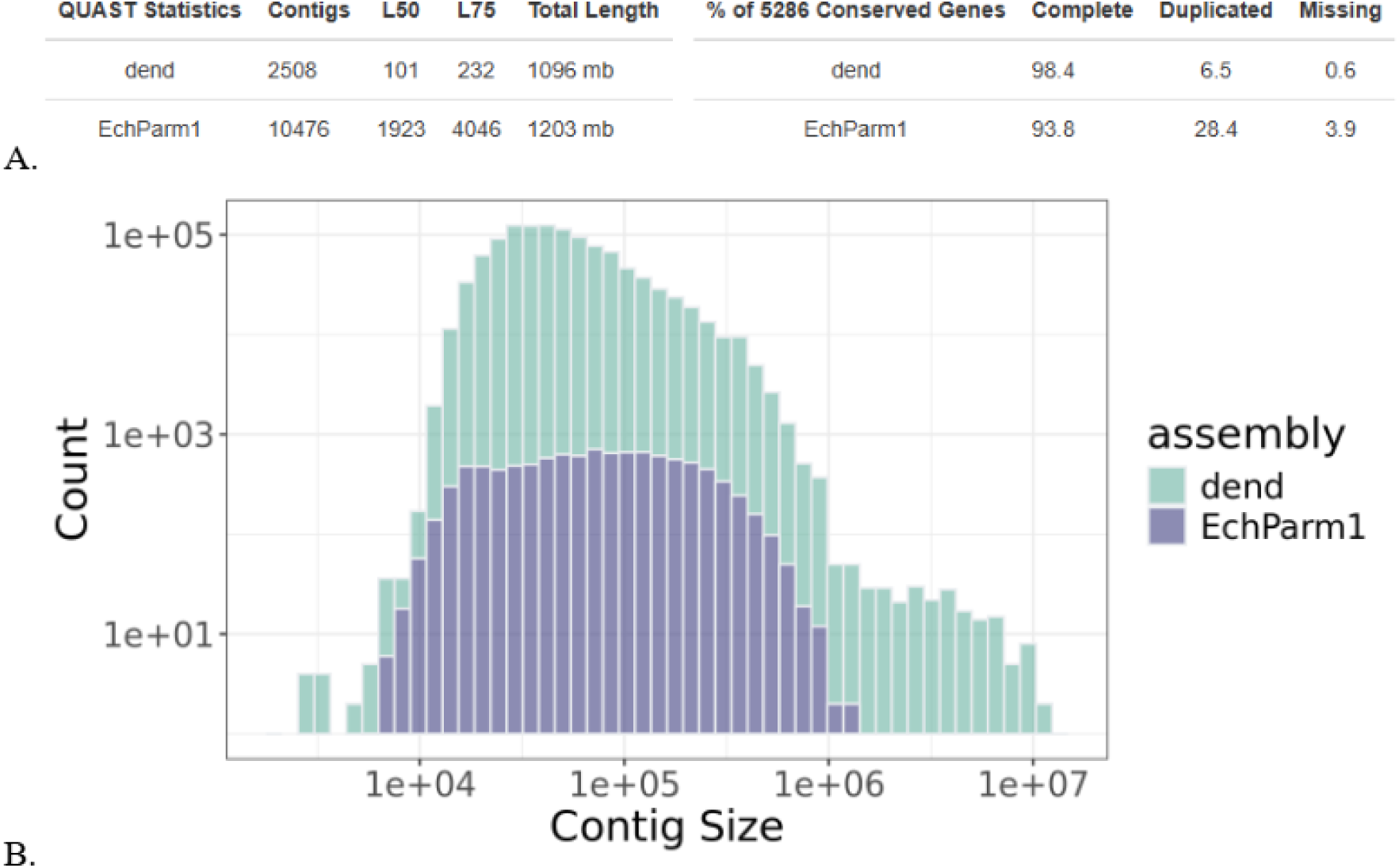
Dendraster Data Assembles with High Completeness and Low Contiguity. a) The EchParm genome, despite being the closest taxonomic reference available, is not closely related nor is it of high quality. Regardless, we can see that the BUSCO completeness for default parameters is already quite good. b) The assembly is in many small pieces, but the largest contigs are the size of small eukaryotic chromosomes.

We can see from the extreme prominence of the first peak in the Genomescope2 output that heterozygous is extremely high in this sample (Figure 9). Relatedly, we see that the assembled genome size is much larger than the total length predicted by Genomescope2. This increased size is caused by diverged haplotypes in a single region assembling as a separate region with high sequence similarity. It is likely that many of the small contigs are actually alternative haplotypes. To address these problems, we can tune the parameters - -purge-max, which controls purging of alternative haplotypes, and -s which controls merging of alternate haplotypes.

**Figure 9:**
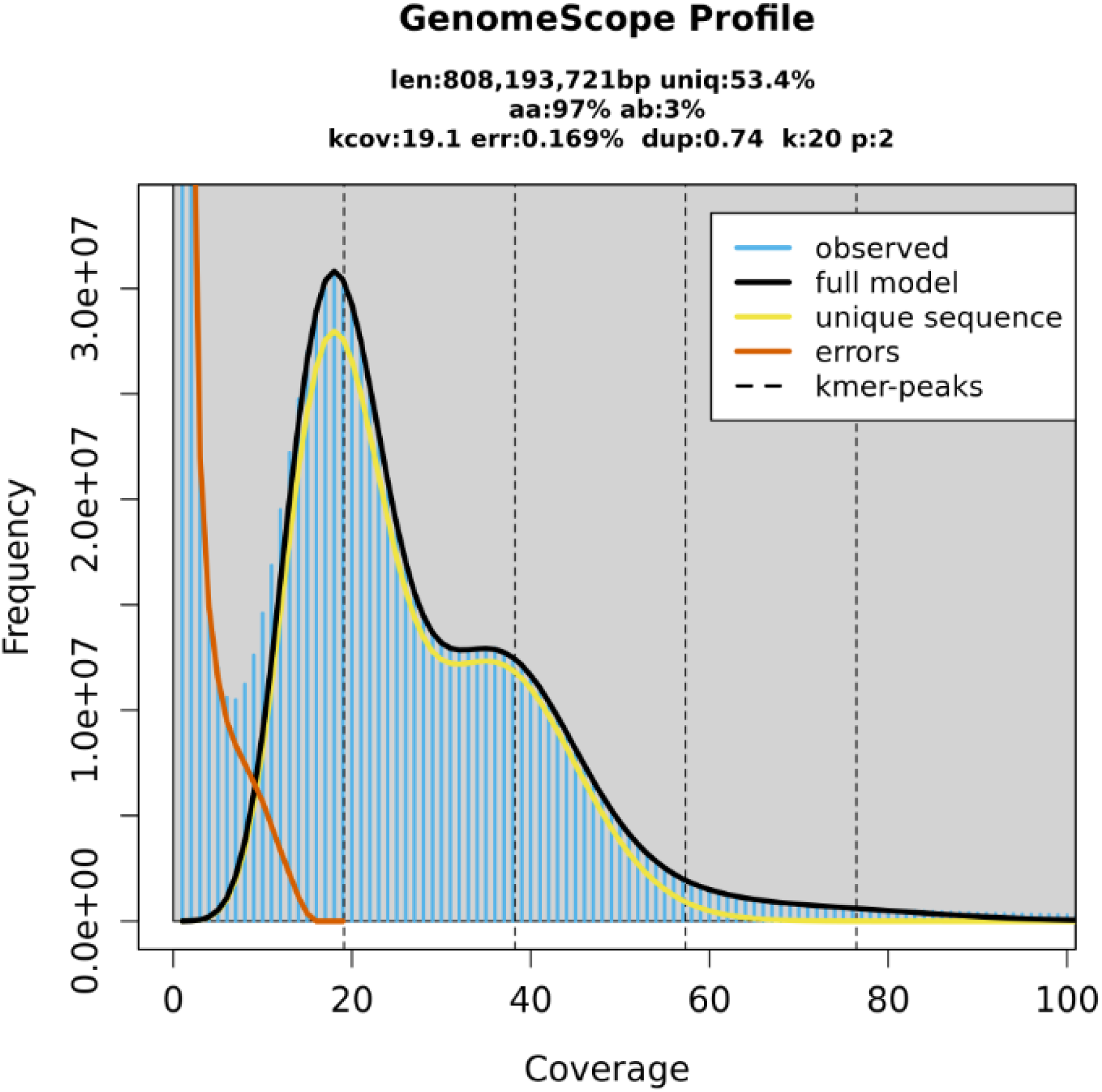
The Genomescope2 kmer Spectrum Reveals High Heterozygosity.

In Figure 10 we see that these changes produce modest improvements. This final assembly, generated de novo from only HiFi reads, is fragmented but highly BUSCO complete and represents a worthwhile addition to genome assembly efforts.

**Figure 10:**
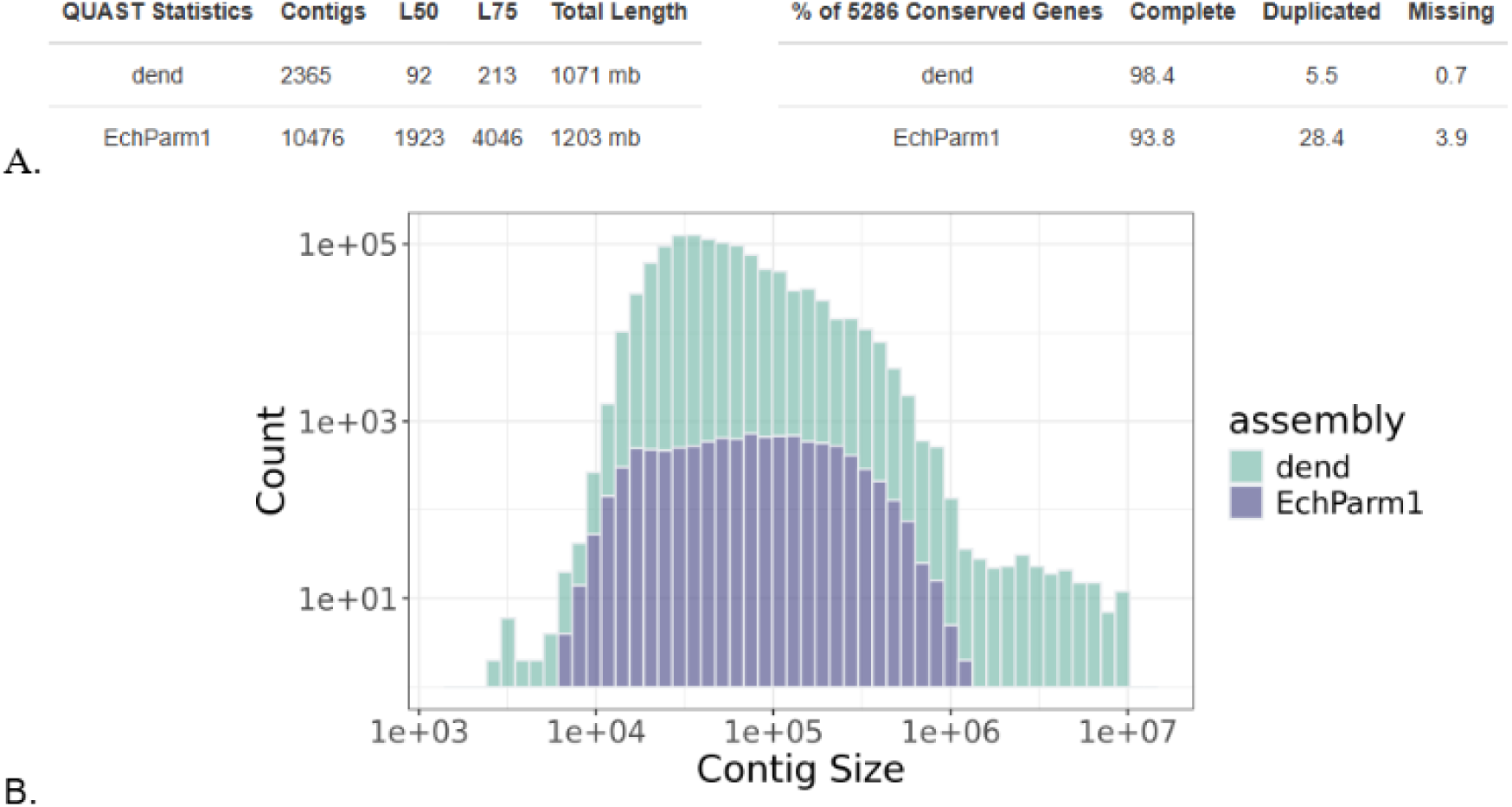
Quick Parameter Tweaks Yield 10% Reduction in L50. a) Modest reductions in overall genome size, contig number, and duplication are achieved without compromising completeness. b) The extreme upper end of contig size is slightly more populated, signaling improvement in contiguity.

### 2.5 Conclusion

When combined with additional data HiFi reads and the HiFiasm assembler can reliably generate chromosome level phased genomes. Without additional data these tools can still generate genomes that meet or exceed the quality of genomes assembled in prior decades with a much lower investment of time and resources. This work provides code and case studies for those interested in getting the best genomes possible from a single HiFi dataset. The visual summary and other outputs from this workflow form a quick and easy basis for choosing assembly parameters. In species like HMR with relevant high quality reference genomes available this approach can yield a more accurate and complete final product by using the reference as a guide for final assembly. When working in clades without existing high quality reference genomes, such as Dendraster excentricus, the approaches detailed here will yield a final product whose broad structural features should be viewed with low confidence. Even so, we show that optimization can yield improvements and that for less studied species HiFi only assembly is still a viable way to improve upon existing genomic resources.

### 2.6 Code Availability

Download and usage information for this project are supported by a public GitHub repository (Pipho, 2026). The repository page displays rich-text documentation which walks users through download, set-up, customization, and implementation. A snapshot of code and documentation at the time of this publication are available through Zenodo at this link: https://doi.org/10.5281/zenodo.18945395

## Acknowledgements

This work was supported by an NIH T32 training grant [KP] and NSF grant 333-2808 [KP, GW]. Heliconius Melpomene sample collection was performed by Joseph Hanly. Dendraster excentricus sample collection was performed by Eric Edsinger. Extraction for this sample was performed by Emma Wallace. Preparation and sequencing for both was conducted at the Duke sequencing core. Many thanks are extended to Avi Heyman, Daniel Levin, Shriya Minocha, Angelina Huang, Qianzi Zhou, Emma Wallace and Chris Shreve for extensive support with design, documentation and user testing.

